# Glycine-Proline driven flexibility and interface-dependent conservation of six-residue Ω-loops in protein complexes

**DOI:** 10.64898/2026.11.05.764545

**Authors:** Surbhi Vilas Tajane, Sanjeevani Pohane, Sucharita Dey

## Abstract

Ω-loops are six- to sixteen-residue, non-regular structural motifs that occur in protein structures, connecting regular regions and forming a well-defined, ordered structural segment. They are functionally important because of their strategic location on the protein surface, where they are often associated with molecular recognition, ligand binding, and catalytic activity. Their non-regular nature enables them to adopt different conformations, which makes them important in protein dynamics and allosteric communication. In this work, we have curated six-residue Ω-loops from diverse protein-protein complexes of varying stability and found them to be abundant across all. We identified 1205 hydrogen-bonded and 6518 non-hydrogen-bonded Ω-loops, leading to an average of ∼1.7 Ω-loops per structure. Glycine and proline were the most enriched residues in the loop, with proline preferentially positioned at the fourth position and glycine distributed across multiple positions, supporting their roles in chain reversal and conformational flexibility. Using ProteinMPNN-based sequence design and Rate4Site evolutionary rate analysis, we show that Ω-loops occupy a relatively constrained sequence space, with interface loops in obligate homodimers and heterodimers more conserved than non-interface loops; the opposite trend was observed in transient and crystal dimers. Mapping known pathogenic and benign mutations onto the Ω-loop present in human proteins, together with a case study of the TEM1 β-lactamase Ω-loop, further illustrates how small sequence changes within these loops can produce crucial structural and functional consequences, leading to altered enzyme activity and antibiotic resistance. Altogether, we report that Ω-loops are not merely connectors of secondary structure but are structurally constrained, evolutionarily significant elements that contribute to protein folding, stability and function.

## Introduction

Proteins are fundamental macromolecules responsible for every cellular function, including catalysis, structural support, signalling, and regulation.^1^ Despite vast diversity, proteins often consist of particular repeating stretches of sequences or structural elements that are crucial to structure-function. These are referred to as protein ‘motifs’. Generally, motifs are distributed across the protein; however, studies have shown that irregular secondary structures more often contain repeating motifs.^2,3^ These repeats, either in the form of sequence or structural motifs, contribute to stability and also play specific functional roles. The sequence motifs are short, functionally important, and evolutionarily conserved amino acid patterns ranging from 3 to 13 residues. They do not adopt any specific secondary structure, but they have strong evolutionary recognition features.^4^ For example, the GXXXG and AXXXA motifs are among the most highly enriched five-residue sequence patterns that facilitate α-helical packing in both membrane and globular proteins.^5,6^ Apart, there are structural motifs, which consist largely of loops and turns, and make up over 30% of globular proteins. These motifs play a vital role in protein stability by enabling a variety of noncovalent interactions and supporting the arrangement of higher-order secondary structures. They are also essential for reversing the direction of the polypeptide chain and linking regular structural elements such as α-helices and β-sheets. β-turns and γ-turns are most common, along with α-turns less frequently observed.^3,7,8,9^ Despite their prevalence, these motifs are more challenging to classify than α-helices or β-sheets because of their structural diversity, and they typically occur in regions where the polypeptide chain changes direction.^10^ They help compact globular packing and protein folding pathways.^11^ So, these structural motifs, either exposed to the surface and important for molecular recognition, or at the interface helpful for protein flexibility, and at the core, are helpful for tight packing and structural stability.^12,13,14, 15^

Other short nonregular structural elements include β-hairpin turns and omega (Ω) loops. The β-hairpin turns are three to five residues long and more easily categorized than the longer segments of nonregular secondary structure.^16–18^ Whereas Ω loops are longer (6 to 16 residues), yet, well-defined structural motifs, first defined by Leszczynski JF and Rose GD in 1986.^2,19,20^ This structural motif helps reverse the direction of the polypeptide chain in three-dimensional space, such that two terminal residues close to each other form a Greek letter omega (Ω).^20^ To distinguish Ω-loops from β-hairpins, γ-turns, and so on, Ω-loops are typically defined as polypeptide segments of about 6 to 16 residues with a distance constraint of less than 10 Å between their terminal residues. The distance must be under two-thirds of the greatest internal distance within that fragment.^19,20^ Unlike α-helices or β-sheets, Ω-loops do not exhibit repetitive backbone dihedral angles or regular hydrogen bonding patterns, but Ω-loops are compact, well-defined non-regular motifs, not just disordered linkers^2,19,20^, and often play crucial role in protein function. Ω-loops are often located at the protein surface where they can contribute to substrate or inhibitor binding, molecular recognition, organization of catalytic sites, and protein-protein interactions.^2,19^ More generally, loop flexibility also plays an important role in protein conformational dynamics and allosteric communication.^21^ In some proteins, they have also been proposed to function as independent structural or evolutionary modules, such as changes in loop composition can lead to functional changes in the protein.^22,23,24^

Although Ω-loops are frequently associated with molecular recognition and functional plasticity, they can also contribute to protein stability. Their stabilizing effect may arise through hydrogen bonding, favourable packing interactions, and their role in linking adjacent regular secondary-structural elements.^2^ Thus, the Ω-loop is important not only as a functional surface motif but also as a structural determinant of protein complex stability. Its functional importance is illustrated by several well-known examples, such as the replacement of a surface loop in ribonuclease A, which conferred angiogenic activity and altered substrate specificity.^24^ Likewise, in Gram-negative bacteria, the E. coli TEM1 beta-lactamase enzyme, a 16-residue Ω-loop is present close to the active site, and the enzyme is found to hydrolyze penicillin and early cephalosporins but is unable to hydrolyze oxyimino cephalosporins due to their large side chains. However, a triple mutant at the Ω-loop, W165Y/E166Y/P167G, increases the Ω-loop volume owing to a large conformational change, allowing oxyimino cephalosporins (with bulky side chains) to fit within it and hydrolyze it.^25^ So, the Ω-loops may fall under the category of functional loops.

In the current study, we investigated the abundance and role of Ω-loops across a large, nonredundant dataset comprising 26152 protein-protein complexes that differ in oligomeric state and stability, and assessed their contributions to the interface. We classified the identified Ω-loops based on their spatial location, hydrogen-bonding characteristics, and likely structural/functional role. Further, we characterized them in terms of their geometry and size.

## Methods

### Dataset curation

We used the same curated dataset as in our earlier study on the abundance of AXXXA motifs.^6,26^ Our non-redundant dataset consists of 6792 Obligate Homodimers (OD), 1359 Heterodimers (HetD), 12378 Monomers, 4923 Crystal Dimers (CD), curated using 3Dcomplex^27^ and QSbio.^28^ The most accurate biological assemblies (X-ray structures) of proteins from QSBio annotations and the structure data were taken from the Protein Data Bank (PDB).^29^ The structures represent a non-redundant set filtered at 90% sequence identity and annotated as physiological assemblies with an error rate below 5%, as obtained from QSbio. We also used a separate non-redundant set of 700 Transient dimers (TD) from the Benchmark dataset, which are most likely transient in nature.^30^

### Definition and Identification of Ω-loops

We identified candidate 6-16 residue Ω-loops using an in-house Python program. We consider Ω-loops of length L, at least L-1 of which are designated as having a secondary structure of “S’ (bend) or “T’ (turn) using the DSSP program.^31^ To ensure that these stretches resembled the classical Ω-loop geometry, we measured the distance between terminal residues and retained only those loops in which this distance is less than 10 Å, which is 2/3^rd^ of the longest c_α_-c_α_ distance within the fragment.^20^ These segments are classified as ‘Ω’ loops.

### Classification of Ω-loops based on hydrogen bonding

Hydrogen-bonded Ω-loops were identified using the HBPLUS program. Loops in which terminal residues (example, i–i+5 residues in 6 residue Ω-loops) form the hydrogen bond were classified as hydrogen-bonded Ω-loops. Loops lacking a terminal residue hydrogen bond are classified as non-hydrogen-bonded Ω-loops.

### Identification of Ω-loops at the interface

The protein-protein interface was defined as the region that is buried upon complex formation, called the interface or buried surface, and its contributing area is called the buried surface area (BSA). Solvent-accessible surface area (SASA) was calculated using the NACCESS program.^32^ Interface regions were identified using the previously described methodology.^33^ Ω-loops were classified as interface and non-interface loops based on their location. If at least five of the six residues of an Ω-loop are located within the interface region, the loop is classified as an interface Ω-loop. When Ω-loops from both chains were present at the interface, the pair was classified as an interface Ω-loop pair.

### Size and Geometry of the Ω-loops

We have calculated several geometrical parameters for each identified Ω-loop.

i. Size parameters

a. Radius: The radius of the Ω-loop is calculated by first identifying the midpoint between the C*α* atoms of the terminal residues of the loop. Then, the distance from this midpoint to the farthest C*α* atom was taken as the diameter of the Ω-loop. The radius is defined as half of diameter distance^20^
b. Surface area: The surface area of the Ω-loop is calculated using formula 4*Πr*2.
c. Volume: The volume of Ω-loop was calculated using formula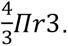
ii. Distance parameters Minimum and maximum residue distances: We calculated the minimum and maximum c_α-_c_α_ distances between two residues c_α_ using the Euclidean distance formula, excluding directly adjacent residues.
iii. Exposure Index: The exposure index is calculated using the following formula: 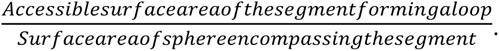^20^

### Characteristics of the interface Ω-loops

We characterized the interface Ω*-*loops in terms of their contribution to buried surface area (BSA) and the loops residue composition -

a. Contribution of individual Ω*-*loops to the interface(%): The area occupied by an individual Ω-loop in the interface region was calculated as the ratio of the interface area contributed by the Ω-loop to the total interface area contributed by that subunit reported in Å², and for the percentage contribution, the value was multiplied by 100.
b. Contribution of individual Ω-loop pairs to the interface(%): The area occupied by Ω-loop pairs in the interface region (Å²) was calculated as the sum of their interface areas, divided by the total interface area of the protein, multiplied by 100.
c. Distance between interface Ω-loop pairs: The distance between Ω-loop pairs at the interface region (Å) was calculated as the distance between centroids, determined using the center of mass (COM).

### Presence of non-protein molecules within Ω-loops

We determined whether any non-protein molecules were present within the identified Ω-loop. For each Ω-loop, COM was calculated, and the distance between the loop COM and each atom of a ligand or ion was measured. If this distance were less than the radius of the Ω-loop, the ligand or ion would be present within the Ω-loop.

### Amino acid propensity of the Ω-loops

Amino acid propensities in Ω-loops were calculated using the following formula:

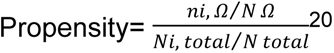

Where,

n_i,Ω **is,**_ The specific amino acid residue, i found in Ω-loops.

N_Ω_, The number of total amino acids found in Ω-loops.

N_i, total_ The specific amino acid residue, i found in total protein.

N_total_, The number of total amino acids found in total protein.

We also visualized the amino acid preference in the Ω-loop using WebLogo.^34^

### Analysis of secondary structures connected by Ω-loops

To analyze the secondary-structure segments connected by Ω-loops, we examined the five residues preceding and five residues following each Ω-loop. Secondary structural information were obtained using DSSP.^31^ A segment of five residue before and after Ω-loop was considered helical if it contained at least 4 out of 5 consecutive H/G/I residues (‘H’ for α-helix, ‘G’ for 3^10^-helix, and ‘I’ for π-helix). Similarly, the presence of at least 4 out of 5 residues were assigned to B/E was considered a β-strand segment. All other cases, including S/T notations (turns/bend), mixed combinations of H/G/I/B/E/S/T, or blank notations are classified as non-regular.^6,26^

### Mutation Data

To evaluate the functional significance of the Ω-loop, we identified already known benign and pathogenic mutations that are present in the Ω loop. Mutation data were obtained from HuVarBase^35^ and curated across the entire human proteome (Acharya et al, unpublished), and these mutations were mapped onto our dataset.

### Conservation and designability of Ω-loops

We analyzed sequence conservation in Ω-loops. To evaluate the evolutionary conservation, we performed Ω-loop sequence similarity searches using BLAST^36^ against the UniRef50^37^ database. Protein sequences containing Ω-loops were used as queries, with selection criteria set to 40-98% sequence identity and 80-100% coverage. Only protein sequences with at least five homologs were retained for further analysis. Multiple sequence alignment (MSA) was generated using MUSCLE(v3.8.1551)^38^, and the resulting alignment was then analyzed using Rate4Site^39^ to calculate residue-wise conservation scores. Rate4Site scores for Ω-loop residues were assessed to evaluate their conservation patterns. In addition, Ω-loops were categorized as interface and non-interface loops, allowing comparison of conservation between these two groups.

The normalized Rate4Site value for each residue in an Ω-loop was calculated by subtracting the most negative Rate4Site value in the chain. For each six-residue Ω-loop, the six normalized Rate4Site values were summed to obtain a loop-level conservation score. In addition, the mean normalized Rate4Site score of each loop was calculated as the sum of the six normalized Rate4Site values divided by six(the loop length). The corresponding chain conservation score was calculated as the mean normalized Rate4Site score for residues present in that chain containing Ω-loop, calculated by the sum of normalized Rate4Site scores for all residues in the chain divided by the number of residues present in that chain. To check the relative conservation of Ω-loops compared with their corresponding protein chains, ΔR4S was calculated for each Ω-loop as the Ω-loop mean R4S score minus the corresponding chain mean R4S score. Negative ΔR4S values indicate greater conservation of the Ω-loop relative to the corresponding chain, whereas positive values indicate lower conservation. The mean ΔR4S was then calculated separately for interface and non-interface Ω-loops within each dataset.

### Designability analysis of Ω-loops

To assess the sequence variability or designability of Ω-loops, we used vanilla ProteinMPNN model(v_48_020)^40^ with a constrained design strategy in which only Ω-loop residues were allowed to vary while the rest of the protein structure was kept fixed. ProteinMPNN was run with a sequence sampling parameter of 12, which often generates duplicate sequences. After removing redundant sequences, unique Ω-loop sequences were considered for further analysis. These filtered sequences were then evaluated using the BLOSUM62 substitution matrix to quantify sequence variability within the Ω-loop regions.

### Classification of Ω-loops

We classified Ω-loops according to their contribution to the stability of the structure or whether they are involved in folding or if they are directly/indirectly associated with a particular function. For stability-associated Ω-loops, intra-loop hydrogen bonds were identified within the Ω-loop using HBPLUS,^41^ excluding interactions between consecutive residues. Both main-chain and side-chain hydrogen bonds were considered. Ω-loops containing four or more intra-loop hydrogen bonds, excluding interaction between consecutive residues, were classified as stability-associated Ω-loops. For folding or secondary-structure connecting Ω-loops, the structural context of Ω-loops was assessed using secondary structure obtained from DSSP. Ω-loops were classified as involved in folding if at least five residues upstream and downstream adopted stable, regular secondary structures, such as α-helices or β-strands. For function or ligand or ion associated Ω-loops were classified as functional loops if they were associated with ligands or ions located within the loop. This was determined by calculating the distance between the center of mass(COM) of the loop and the ligand, if this distance was less than the loop radius, the ligand or ion was considered to be present within the Ω-loop.

## Results and Discussion

### Ω-loops are abundant in obligate protein homodimers

We defined Ω-loops motivated by the criteria used previously by Dasgupta group.^20^ Our definition slightly differs: we have considered Ω-loops of length L, at least L-1 of which are designated as having a secondary structure of “S’ (bend) or “T’ (turn) using the DSSP program (Figure 1).^31^ Ω-loops ranging from 6 to 16 residues were considered, although loops longer than 9 residues are rare. A similar trend was observed in Pal and Dasgupta et al^20^, where the number of Ω-loops decreased with increasing loop length. Hence, we primarily focused on characterizing the six-residue Ω-loops – the most abundant ones, and refer to them as Ω-loops all throughout, although we have provided the characterization data for 7-9 residue loops in the Supplementary information.

**Figure 1:**
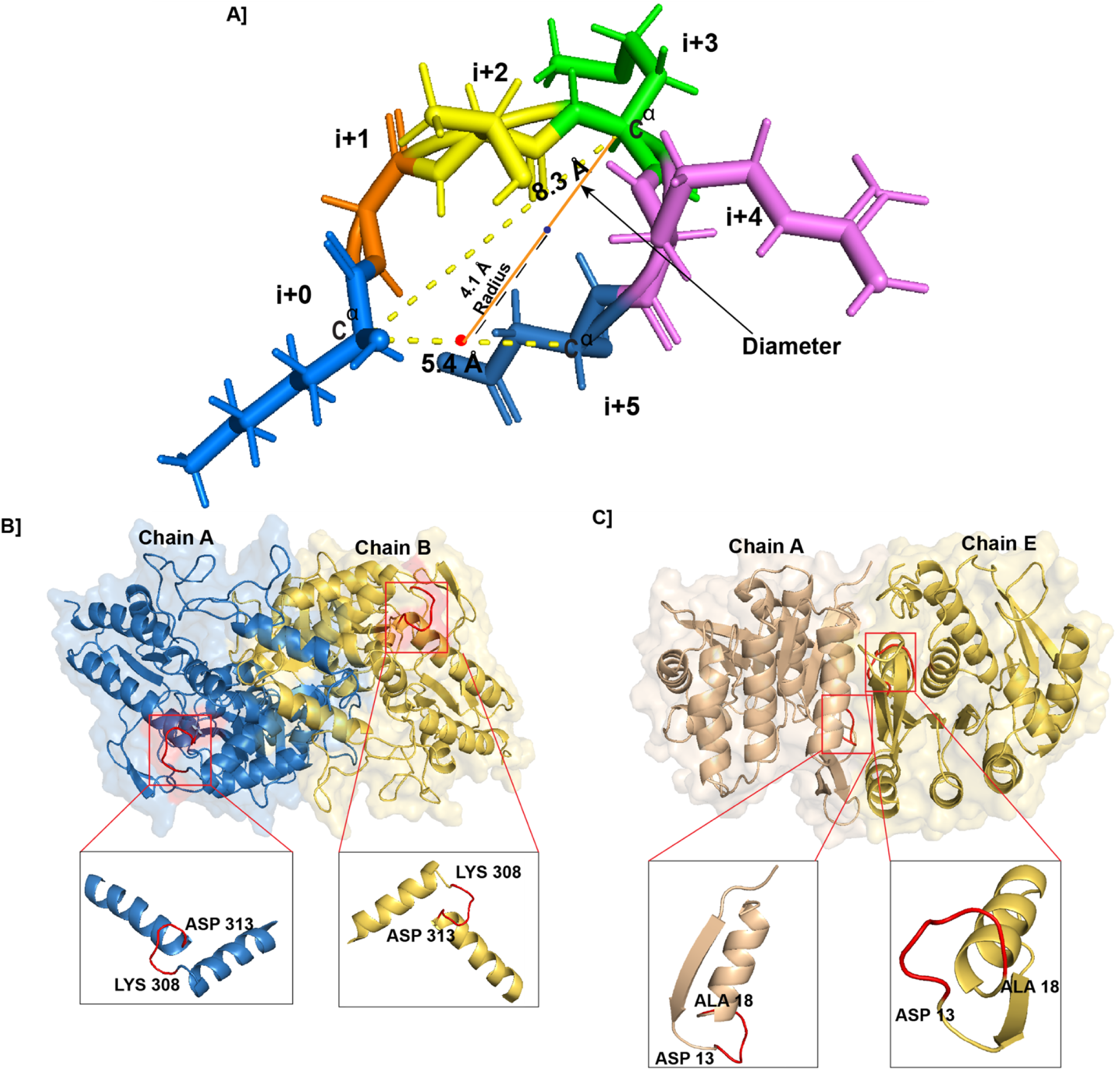
Definition of Ω-loop, Ω-loop pairs, and Ω-loops at protein interfaces. (A) Example of a dimeric protein, alliinase from *Allium sativum* (PDB ID: 2hox), showing an Ω-loop (shown in red colour) spanning residues 308-313. (B) Example of Ω-loop pairs located on the protein surface in the homodimeric protein UDP-N-acetylglucosamine 2-epimerase from *Thermus thermophilus* (PDB ID: 1v4v). Ω-loops are present in both chain A and chain B, corresponding to residues 308-313. (C) Example of Ω-loop pairs located at the protein-protein interface in the homodimeric ADP-bound structure from *Bacillus subtilis* (PDB ID: 2i5b). Ω-loops are present in both chain A and chain E, corresponding to residues 13-18.

We classified Ω-loops into hydrogen-bonded Ω-loops (HBΩ) and non-hydrogen-bonded Ω-loops (nHBΩ) – loops in which the terminal residue formed a hydrogen bond were classified as HBΩ, whereas those lacking such terminal hydrogen bonding were classified as nHBΩ. Across all datasets, we identified 1205 HBΩ-loops and 6518 nHBΩ-loops, leading to an average of ∼1.7 Ω-loops per structure. However, the count of these loops is found to be the least in the case of heterodimers and monomers (∼1 HBΩ and ∼1.4 nHBΩ on average per structure in heterodimers and monomers), while in obligate homodimers it is 1.8 HBΩ and 2.1 nHBΩ and in Transient dimer, 1.9 HBΩ and 2.2 nHBΩ. Overall, nHBΩ-loops were more common than HBΩ-loops across all types of complexes (Table 1).

**Table 1.**
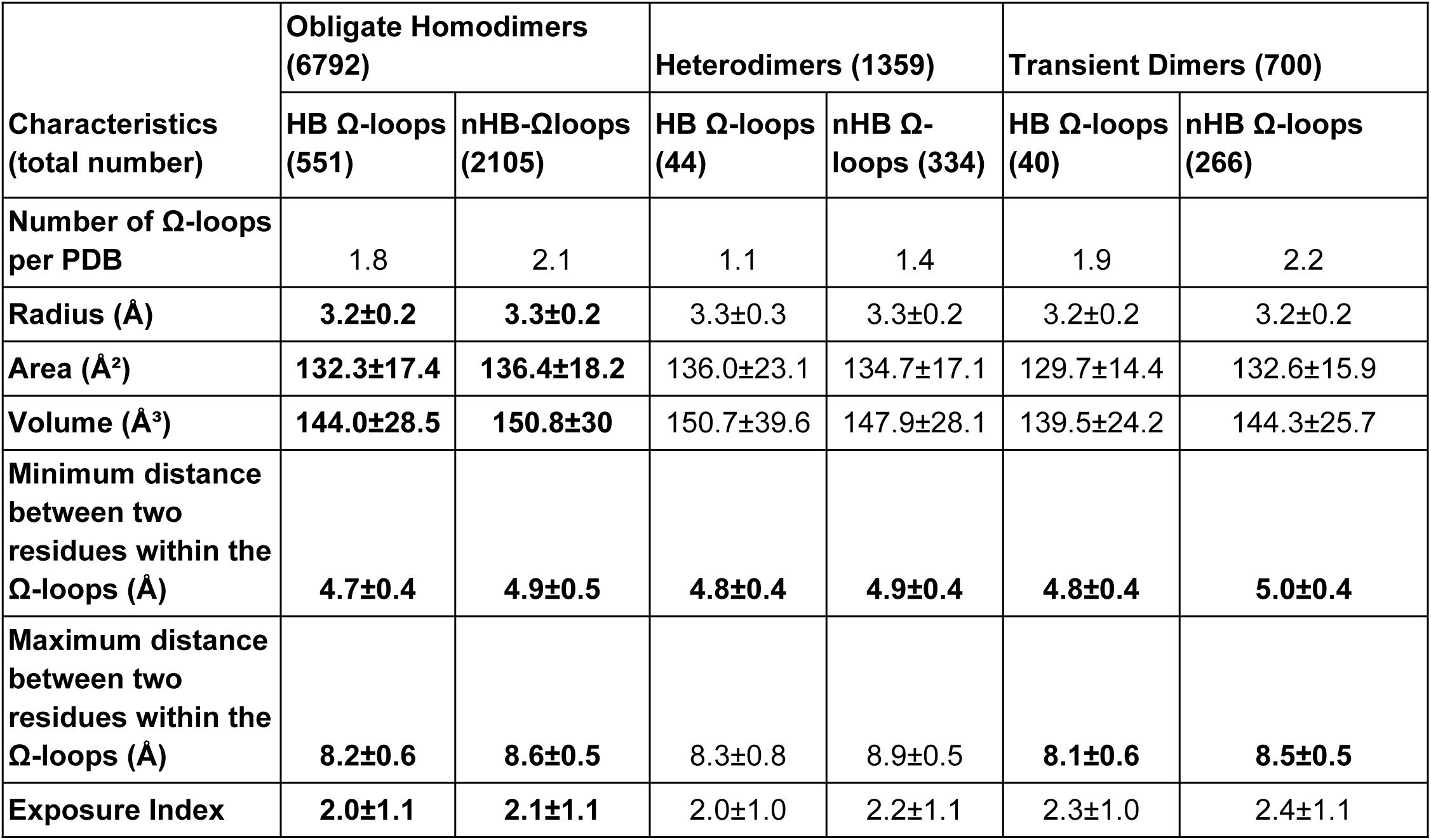
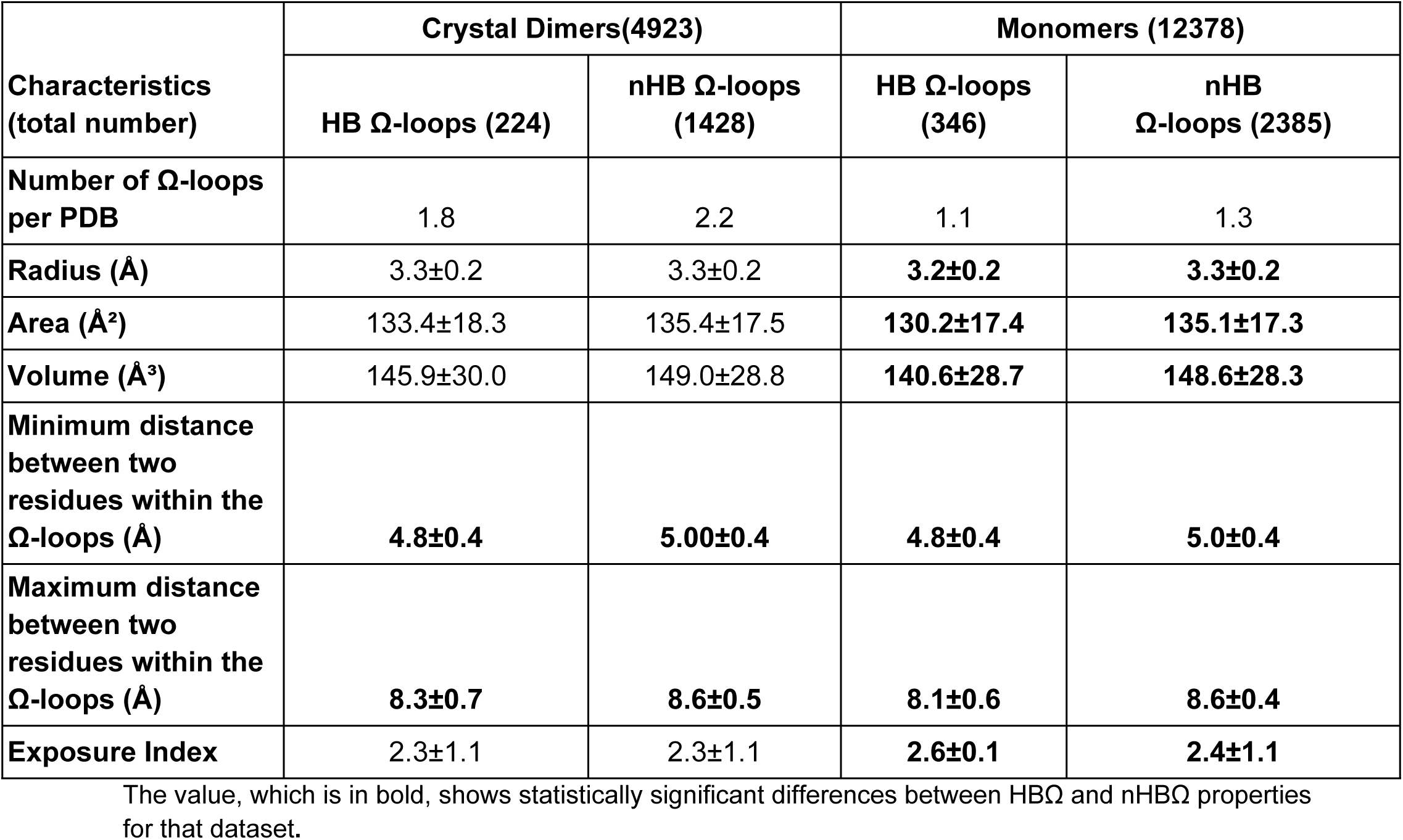
General characteristics of six-residue Ω-loops across different protein-protein complexes.

| **Characteristics (total number)** | **Obligate Homodimers (6792)** |  | **Heterodimers (1359)** |  | **Transient Dimers (700)** |  |
|---|---|---|---|---|---|---|
|  | **HB Ω-loops (551)** | **nHB-Ωloops (2105)** | **HB Ω-loops (44)** | **nHB Ω-loops (334)** | **HB Ω-loops (40)** | **nHB Ω-loops (266)** |
| **Number of Ω-loops per PDB** | 1.8 | 2.1 | 1.1 | 1.4 | 1.9 | 2.2 |
| **Radius (Å)** | **3.2±0.2** | **3.3±0.2** | 3.3±0.3 | 3.3±0.2 | 3.2±0.2 | 3.2±0.2 |
| **Area (Å²)** | **132.3±17.4** | **136.4±18.2** | 136.0±23.1 | 134.7±17.1 | 129.7±14.4 | 132.6±15.9 |
| **Volume (Å³)** | **144.0±28.5** | **150.8±30** | 150.7±39.6 | 147.9±28.1 | 139.5±24.2 | 144.3±25.7 |
| **Minimum distance between two residues within the Ω-loops (Å)** | **4.7±0.4** | **4.9±0.5** | **4.8±0.4** | **4.9±0.4** | **4.8±0.4** | **5.0±0.4** |
| **Maximum distance between two residues within the Ω-loops (Å)** | **8.2±0.6** | **8.6±0.5** | 8.3±0.8 | 8.9±0.5 | **8.1±0.6** | **8.5±0.5** |
| **Exposure Index** | **2.0±1.1** | **2.1±1.1** | 2.0±1.0 | 2.2±1.1 | 2.3±1.0 | 2.4±1.1 |
| **Characteristics (total number)** | **Crystal Dimers(4923)** |  | **Monomers (12378)** |  |  |  |
|  | **HB Ω-loops (224)** | **nHB Ω-loops (1428)** | **HB Ω-loops (346)** | **nHB<br>Ω-loops (2385)** |  |  |
| **Number of Ω-loops per PDB** | 1.8 | 2.2 | 1.1 | 1.3 |  |  |
| **Radius (Å)** | 3.3±0.2 | 3.3±0.2 | **3.2±0.2** | **3.3±0.2** |  |  |
| **Area (Å²)** | 133.4±18.3 | 135.4±17.5 | **130.2±17.4** | **135.1±17.3** |  |  |
| **Volume (Å³)** | 145.9±30.0 | 149.0±28.8 | **140.6±28.7** | **148.6±28.3** |  |  |
| **Minimum distance between two residues within the Ω-loops (Å)** | **4.8±0.4** | **5.00±0.4** | **4.8±0.4** | **5.0±0.4** |  |  |
| **Maximum distance between two residues within the Ω-loops (Å)** | **8.3±0.7** | **8.6±0.5** | **8.1±0.6** | **8.6±0.4** |  |  |
| **Exposure Index** | 2.3±1.1 | 2.3±1.1 | **2.6±0.1** | **2.4±1.1** |  |  |

The value, which is in bold, shows statistically significant differences between HBΩ and nHBΩ properties for that dataset.

### Characteristic features of Ω-loops across datasets

We evaluated the structural characteristics of HBΩ and nHBΩ loops across five different protein datasets: Obligate Homodimers (OD), Heterodimers (HetD), Transient Dimers (TD), Crystal Dimers (CD), and Monomers. For all identified loops, we calculated radius, area, volume, maximum and minimum intra-loop distances, and exposure index. The distinct and highlighted contrasts between HB and nHB Ω-loops in each set are outlined. (Table 1) (Supplementary Tables 1 and 2)

**Table 2.** General Characteristics of six-residue Ω-loops present at the interface in Obligate Homodimers, Heterodimers, Transient Dimers, and Crystal Dimers.

| **Characteristics** | **Obligate Homodimers** |  | **Heterodimers** |  | **Transient Dimer** |  | **Crystal Dimers** |  |
|---|---|---|---|---|---|---|---|---|
|  | **HB Ω-loops (43)** | **nHB Ω-loops (181)** | **HB Ω-loops (0)** | **nHB Ω-loops (15)** | **HB Ω-loops (4)** | **nHB Ω-loops (6)** | **HB Ω-loops (6)** | **nHB Ω-loops (57)** |
| **Number of loops per PDB** | 1.7 | 1.9 | - | 1 | 2 | 1.5 | 1.5 | 1.6 |
| **<Area contribution by Ω-loops to BSA (Å<sup>2</sup>)> (% contribution to BSA)** | 217.9± 78.7 (7.8 ± 4.9%) | 214.9 ± 74.2 (7.9 ± 5.0 %) | - | 212.7 ± 132.2 (14.1± 10.3 %) | 158.3± 4.8 (13.2 ± 1.9%) | 285.2 ± 68.7 (26.2 ± 6.8%) | 196.3± 140.3 (24.7 ± 23.1%) | 207.1± 80.1 (19.0 ± 11.4%) |
| **Ω-loop Pairs** |  |  |  |  |  |  |  |  |
|  | **16 PAIRS** | **65 PAIRS** | **-** | **-** | **2 PAIRS** | **2 PAIRS** | **2 PAIRS** | **19 PAIRS** |
| **<Area contribution by Ω-loops pairs to BSA (Å<sup>2</sup>)> (% contribution to BSA)** | 418.2± 152.1 (6.8 ± 4.6 %) | 445.7± 151.9 (8.0 ± 5.2 %) | - | - | 316.7± 8.6 (13.2 ± 2.4 %) | 628.6± 54.8 (30.4 ± 0.6 %) | 405.4± 343.6 (31.5 ± 31.4 %) | 400.7± 151.9 (19.3 ± 11.4 %) |
| **Distance Between Ω-loop pairs (COM) found at interface region (Å)** | 28.5 ± 11.2 | 26.2± 12.4 | - | - | 26.8 ± 15.9 | 20.3 ± 8.5 | 21.3 ± 14.5 | 18.5 ± 8.7 |

HB: Hydrogen bonded; BSA: Buried surface Area; COM: Center of mass

The number of Ω-loops per PDB entry varied among datasets, but in all cases, nHBΩ-loops were more prevalent than HBΩ-loops. This consistent predominance of nHBΩ-loops across Obligate Homodimers, Heterodimers, Transient Dimers, Crystal Dimers, and Monomers supports earlier observations that Ω-loops generally lack specific hydrogen bonds like other structural motifs.^2,19^ This is also consistent with structural analyses describing Ω-loops as irregular, flexible, and often solvent-exposed regions that typically lack specific hydrogen bonds.^2,19^ The average radius of Ω-loops was similar across all datasets and showed little difference between HBΩ-loops and nHBΩ-loops. This suggests that hydrogen bonding has little influence on the overall radial extent of Ω-loops and that their geometry is governed more by intrinsic loop constraints than by the presence of stabilizing intraloop hydrogen bonding. In contrast, nHB Ω-loops generally showed a slightly larger surface area and volume than HBΩ-loops, indicating HBΩ-loops tend to compact conformations, whereas nHBΩ-loops are relatively more expanded. This indicates that intraloop hydrogen bonds may restrict conformational flexibility, while nHB loops remain structurally broader. The minimum intraloop residue distance differed between HB and nHB Ω-loops. In obligate homodimers, for example, HBΩ-loops showed a shorter minimum distance (4.7 ± 0.4 Å) than nHBΩ-loops (4.9 ± 0.5 Å). A similar trend was observed across Heterodimers, Transient Dimers, Crystal Dimers, and Monomers. Since the shortest distance involved mostly terminal residue Cɑ atom (i+0---i+5), this lower minimum distance in HBΩ-loops may facilitate terminal hydrogen-bond formation. The maximum intra-loop distance also tended to be slightly smaller in HBΩ-loops and nHB loops in several datasets, including in obligate homodimers (8.2 ± 0.6 vs 8.6 ± 0.5 Å), heterodimers (8.3 ± 0.8 vs 8.9 ± 0.47 Å), and monomers (8.1 ± 0.6 vs 8.6 ± 0.4 Å). These trends suggest that hydrogen bonding favors tighter packing of residues within the loop. The exposure index showed that nHBΩ-loops were generally more solvent-exposed than HBΩ-loops (2.1 ± 1.1 vs 2.0 ± 1.1). This suggests that nHB loops are more frequently located at or near protein surfaces, where they may contribute to structural flexibility and facilitate intermolecular interactions. By contrast, HBΩ-loops appear to be somewhat less exposed, indicating that they may occupy more constrained or partially buried environments within protein structures. (Table 1)

### Characteristics of Ω loops located at the protein-protein interface

Obligate homodimers contained the largest number of interface Ω-loops, including 43 HBΩ-loops and 181 nHBΩ-loops, indicating that Ω-loops make a substantial contribution to the interfaces. No HB Ω-loops were observed at heterodimer interfaces, whereas transient dimers and crystal dimers contained only 4 and 6 HB Ω-loops, respectively. In contrast, nHB Ω-loops were present in all dimer classes, with the highest number again observed in obligate homodimers (181), followed by crystal dimers (57), heterodimers (15), and transient dimers (6). The average number of interface Ω-loops per PDB entry showed only minor variation across the different dimer types. The interface area occupied by individual Ω-loops varied across datasets. In obligate homodimers, HBΩ-loops occupied an average of 217.9 ± 78.7 Å², corresponding to 7.8% of the interface, while nHBΩ-loops occupied 214.9 ± 74.2 Å², corresponding to 7.9% of the interface. In heterodimers, nHB Ω-loops showed larger variability in interface area (212.7 ± 132.2 Å²; 14.1%). Transient dimers showed the highest relative contribution from nHB Ω-loops (285.2 ± 68.7 Å²; 26.2%), indicating substantial key participation at these interfaces. In crystal dimers, HB Ω-loops occupied 196.3 Å² (24.7%), whereas nHB Ω-loops occupied 207.1 Å² (19%).(Table 2)

When Ω-loops formed inter-subunit pairs, obligate homodimers again showed the highest numbers (16 HB pairs and 65 nHB pairs). Fewer pairs were observed in transient and crystal dimers (2 HB and 2 nHB in TD; 2 HB and 19 nHB in CD). Interface areas contributed by Ω-loop pairs varied considerably. In obligate homodimers, HB pairs occupied an average of 418.2 ± 152.1 Å², whereas nHB pairs were larger (445.7 ± 151.9 Å²). In crystal dimers, HB (405.4 ± 343.6 Å²) and nHB pairs (400.7 ± 151.9 Å²) show comparable interface area. Transient dimers, in contrast, exhibited the largest nHB pair areas (628.6 ± 54.8 Å²; 30.4%), but very few loop pairs are present. The spatial arrangement of Ω-loop pairs, measured by COM distances, varied across dimer classes. Obligate homodimers generally showed larger inter-loop separations, transient dimers showed intermediate distances, and crystal dimers displayed the shortest separations.(Table 2) We also calculated these properties for Ω-loops of length 7-9 residues. Ω-loops longer than 9 residues were very few, consistent with the earlier observation that longer loops are relatively rare.^20,42^(Supplementary Tables 3-8)

### Loop-mediated interactions

Analysis of loop contents across different oligomeric states showed that most Ω-loops contained only protein atoms located within the Ω-loop COM-radius region, while water, ligands, and ions occurred less frequently. An analysis showed that 72.2%-80.6% of the Ω-loops present in the analysis contained protein atoms alone, predominantly from the main chain rather than the side chain. Water molecules were the most frequently observed non-protein component in all datasets. In monomers, water was present in 26.8% of the loops, whereas ions such as sodium, chloride, and calcium together accounted for 0.5%, and ligands such as FGX, FAD, A52, and IPA accounted for 0.2%. In obligate homodimers, water was present in 26.9% of the loops, followed by ions (0.8% calcium, fluoride, sodium, and potassium) and ligands (0.5% malonate, NAD, phosphate). In heterodimers, water was present in 19.0% of loops, with a small contribution of sodium ions (0.4%). In crystal dimers, water was present in 20.4% of loops, while ligands (SO₄ and citrate, 0.3%) and sodium ions (0.1%) were rarely observed. (Figure 2, Supplementary Table 9)

**Figure 2:**
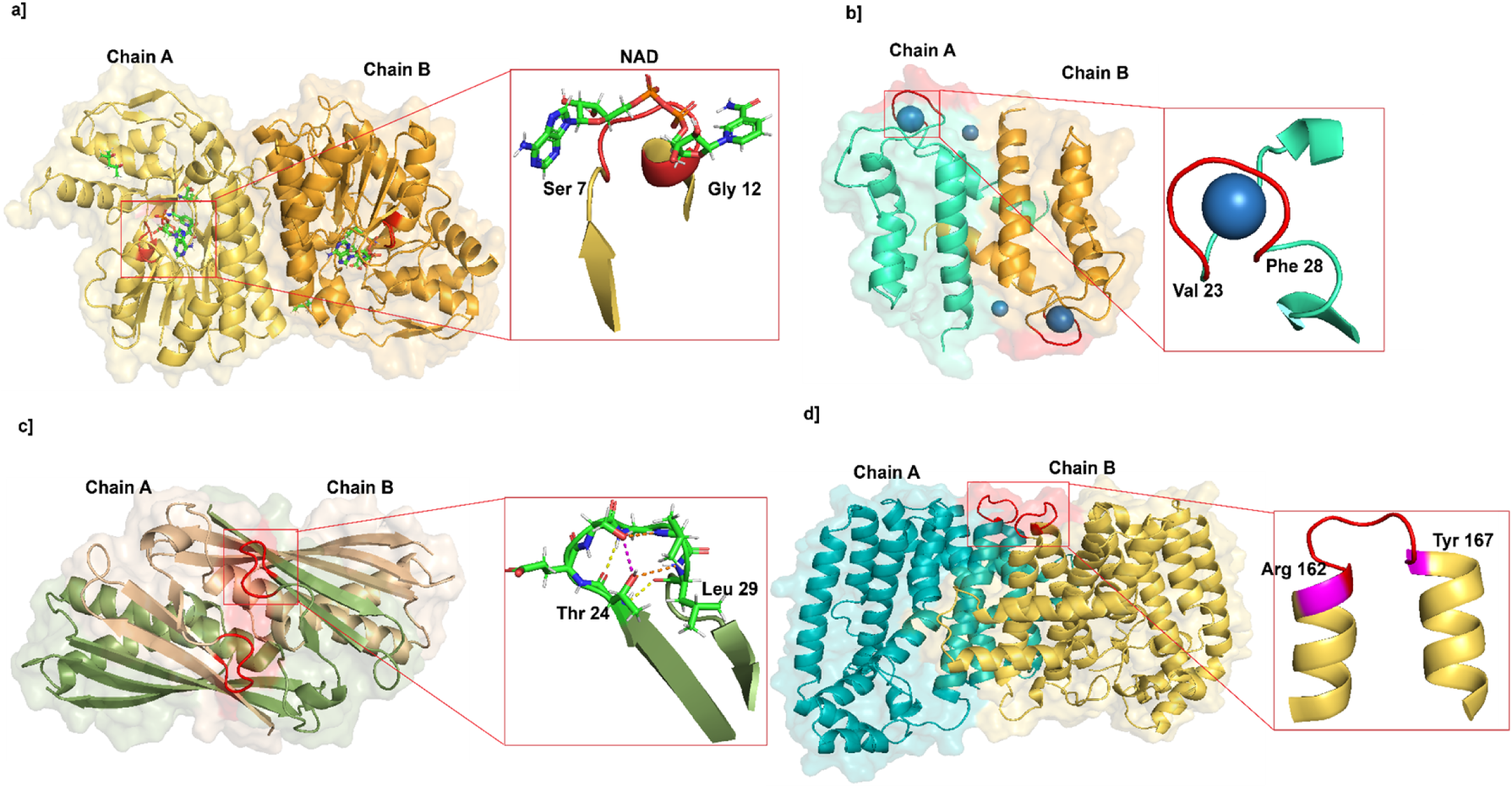
Structural and functional Classification of Ω-loops. (a) Example of a homodimeric protein, L-3-hydroxynorvaline-bound L-threonine dehydrogenase from *Thermoplasma volcanium* (PDB ID: 3ajr), showing an Ω-loop spanning residues 7-12 with the NAD cofactor. (b) Example of Ω-loops in a homodimeric metal-binding protein from *Homo sapiens* (PDB ID: 2wcb). Ω-loops are present in both chain A and chain B, corresponding to residues 23-28, and consist of Na⁺ ion. (c) Example of Ω-loops in the homodimeric protein OsmC from *Escherichia coli* (PDB ID: 1qwi). Ω-loops are present in both chain A and chain B (residues 24-29) and form six hydrogen bonds that contribute to structural stability. (Yellow dashed lines indicate main chain-main chain hydrogen bonds, orange dashed lines indicate main chain-side chain hydrogen bonds, and pink dashed lines indicate side chain-side chain hydrogen bonds.) (d) Example of Ω-loops in the homodimeric chimeric FPP/GFPP synthase from *Artemisia spiciformis* (PDB ID: 4kk2). Ω-loops (residues 162-167) connect regular secondary structure elements (e.g., helices) in both chain A and chain B and may facilitate proper protein folding.

### Amino acid propensity and secondary structure preference

We calculate amino acid preferences and propensities across all datasets, and glycine and proline emerge as the dominant residues in Ω-loops, showing strong positional preferences at multiple sites within the Ω-loop. (Supplementary Figure 1) This trend was also supported by amino acid propensity calculation for monomers, obligate homodimers, heterodimers, transient dimers, and crystal dimers (Supplementary Table 10), in which glycine and proline consistently displayed the highest propensities, confirming their preferential occurrence in loop regions. These observations align with those of Dasgupta et al. (2004), where Gly and Pro were reported as enriched residues in Ω-loops.^20^ Proline showed the strongest preference at the fourth position, consistent with its known role in promoting turn and imposing conformational constraints on the polypeptide backbone. Its propensity values ranged from 1.24 to 1.76, with the highest value observed in obligate homodimers. Glycine, in contrast, was distributed across several positions within the loop, consistent with its role in providing conformational flexibility. Its propensity values ranged from 1.92 to 2.10 across different oligomeric states. Together, these results support that the Ω-loop preferentially accommodates residues that favour local flexibility and chain reversal (Supplementary Figure 2).^19^

In contrast, hydrophobic residues generally showed reduced propensities in Ω-loops. In particular, isoleucine and valine (<0.7) exhibited consistently low values across the dataset, supporting the earlier observation that bulky aliphatic residues are disfavoured in these regions. Also, preferences were observed for negatively charged polar residues such as aspartate and glutamate. Additionally, residues such as serine and threonine also showed comparatively elevated propensities, further supporting previous findings that these residues are also favored in loop regions.

We also checked whether Ω-loops act as connectors between regular secondary-structure elements by analyzing the secondary-structure segments flanking Ω-loops across all datasets. Ω-loops predominantly connect either NR(non-regular)-NR or helix-NR segments, whereas connections between two regular secondary-structure elements were less frequent. (Supplementary Figure 3, Supplementary Table 11)

### Conservation and designability of Ω-loops

To assess the sequence variability and designability of Ω-loops, we applied ProteinMPNN to generate multiple new sequences for each Ω-loop that potentially take the same fold. Although 12 sequences were generated per Ω-loop, only a few Ω-loops yielded fully unique sequences, whereas duplicate sequences were common in most cases, suggesting that Ω-loops occupy a relatively constrained sequence space. This may indicate limited designability.

To quantify sequence variability, we calculated the average BLOSUM62 score using the uniquely designed sequences for each Ω-loop. Most Ω-loops in interface and non-interface regions showed an average BLOSUM62 score of 10-25 across all datasets. In transient dimers and crystal dimers, interface Ω-loops showed, on average, lower BLOSUM62 scores than non-interface Ω-loops, suggesting relatively higher sequence variability at the interfaces. In contrast, non-interface Ω-loops generally showed higher scores, indicating a tendency towards more conservation. Interestingly, an opposite trend was observed in obligate homodimers and heterodimers, where interface Ω-loops showed less variability, hence were more conserved at the interfaces. This reflects a stronger evolutionary constraint imposed on permanent interaction interfaces. Only a small number of Ω-loops fell within the very low score ranges (<0), indicating that most of the designed (new) sequences retain favourable substitutions (Supplementary Figure 4). Overall, these results indicate that Ω-loops exhibit constrained sequence variability, with a tendency toward conserved substitutions that likely help maintain structural integrity.

We also examined the relative evolutionary conservation of Ω-loops using the Rate4Site – a conventional way of evaluating sequence conservation. We compared the mean normalized Rate4Site score of each Ω-loop with the mean score of its corresponding protein chain. Negative ΔR4S values indicate greater conservation of the Ω-loop as compared to the chain, whereas positive values indicate lower conservation. In obligate homodimers, interface Ω-loops showed lower Rate4Site values, showing a mean ΔR4S of -0.273 compared to -0.036 for non-interface Ω-loops, indicating greater conservation at the interface. Similarly, heterodimer interface Ω-loops showed a mean ΔR4S of -0.524 compared to 0.125 for non-interface Ω-loops. In contrast, the transient-dimer interface Ω-loops showed a positive mean ΔR4S 1.117 compared with 0.036 for non-interface Ω-loops, while crystal-dimer interface Ω-loops showed a mean ΔR4S of 0.087 compared with −0.007 for non-interface Ω-loops. Differences between interface and non-interface Ω-loops were significant for obligate homodimers and heterodimers(p < 0.05). Monomer Ω-loops, which lack interface regions, exhibited moderate conservation overall. (Supplementary Table 12)

The ProteinMPNN-based analysis was broadly consistent with the conservation patterns observed with ΔR4S, particularly in transient and crystal dimers, where interface Ω-loops also tended to show lower BLOSUM scores.^43^ To check whether more conserved Ω-loops were more buried at protein-protein interfaces, we analyzed the relationship between the loop interface area and Ω-loop Rate4site conservation score in obligate homodimers. Conserved Ω-loops show more interface area than the least-conserved Ω-loops, consistent with interface area being significantly negatively correlated with the continuous Rate4Site conservation score (Spearman ρ = -0.302, p = 0.000429), indicating that more conserved Ω-loops tend to bury a larger surface area at the protein-protein interface. Also, the solvent-accessible surface area(Spearman ρ = +0.257, p = 0.0029) and exposure index(Spearman ρ = 0.261, p = 0.0025) of those loops were lower in conserved Ω-loops. Conserved interface Ω-loops are therefore more buried against the surface and less exposed to solvent (Supplementary Figure 5). The crystal dimer, heterodimer, and transient dimer datasets contained very few interface Ω-loops to test the relationship.

### Importance and additional role of Ω-loops

We classified Ω-loops across different oligomeric states according to their inferred roles in folding, stability, and function. The total number of Ω-loops was highest in obligate homodimers and monomers compared to other oligomeric states. Among the classified loops, folding-associated Ω-loops were the most abundant across all datasets, followed by stability-associated loops, suggesting that connecting the secondary-structural element is one of the most common roles of Ω-loops. Overlaps between the categories were limited, with few or no Ω-loops shared between folding-functional and stability-functional classes, indicating that Ω-loops rarely perform multiple roles simultaneously. Notably, the majority of Ω-loops were not assigned to any of the three categories, suggesting they may primarily contribute to maintaining the overall structural framework rather than to specific functional roles.(Figure 2) (Table 3)

**Table 3.** Classification of Ω-loops based on their predicted roles in protein folding, structural stability, and function across different oligomeric states.

|  | **Total number of Ω-loops** | **Folding** | **Stability** | **Function** | **Folding & Stability** | **Folding & function** | **Stability & function** | **Not defined** |
|---|---|---|---|---|---|---|---|---|
| **Obligate Homodimer** | 2656 | 365 | 108 | 19 | 23 | 10 | 0 | 2131 |
| **Heterodimer** | 378 | 44 | 11 | 1 | 5 | 0 | 0 | 317 |
| **Transient Dimer** | 306 | 43 | 6 | 0 | 2 | 0 | 0 | 255 |
| **Crystal Dimer** | 1652 | 273 | 84 | 9 | 6 | 0 | 0 | 1280 |
| **Monomer** | 2731 | 440 | 88 | 15 | 38 | 2 | 1 | 2147 |

To assess the structural predictability of Ω-loops and determine whether prediction confidence varies with conservation, we modelled 77 representative structures containing 77 more- and less-conserved Ω-loops using ColabFold (AlphaFold2-multimer v3). From each dataset, up to five more conserved and less conserved Ω-loops were selected, separately for interface and non-interface loops, fewer loops present in the heterodimer and transient dimer datasets, resulting in 77 Ω-loops overall (Supplementary Table 13). For each Ω-loop, the loop pLDDT score was calculated as the mean pLDDT value of the six residues comprising the Ω-loop. Most Ω-loops were predicted with high confidence, with 42 of 77 loops(55%) showing pLDDT values >90 and 65 of 77 loops (84%) showing pLDDT >70. Only 12 loops (16%) had pLDDT <70. Importantly, the prediction confidence was not associated with conservation, conserved Ω-loops mean pLDDT 86.5 compared with 82.1 for least-conserved Ω-loops (Welch’s t-test, p = 0.201), and in continuous Rate4Site conservation score was not significantly correlated with loop pLDDT (Spearman ρ = -0.125, p =0.28). These results indicate that both conserved and least-conserved Ω-loops can generally be modelled with high confidence, thus evolutionary conservation does not appear to be a major determinant of Ω-loop prediction confidence in the present dataset.

### Mutation mapping within Ω-loops

To examine the potential functional significance of the Ω-loop in addition to their structural role, we mapped already known benign and pathogenic mutations onto Ω-loop-containing proteins. Benign and pathogenic mutations from Huvarbase^35^ were curated for the entire human proteome (Acharya et al, unpublished), and we mapped them onto our dataset of obligate homodimers, monomers, and heterodimers. We identified 13 heterodimers, 19 obligate homodimers, and 93 monomers of human proteins in our dataset that had benign, pathogenic, or both types of mutations. For example, in the homodimeric human glutathione S-transferase structure (PDB ID: 1XW6), an Ω-loop spanning residues 146-151 was identified. A mutation at residue 151, N151D, has been reported as associated with breast cancer.^44^

As a case study, we looked into a well-known example, TEM-1 β-lactamase, in which a 16-residue Ω-loop (residues 164-179) is present in the active site. The W165Y/E166Y/P167G triple mutant induced a larger conformational change in this loop, expanding the cavity and shifting substrate preference from penicillins and early cephalosporins towards the bulky oxyiminocephalosporin and ceftazidime.^25^ and consistent with this, we calculated Ω-loop volume, which showed an increase from 1365.3 Å³ in the wild-type structure to 1687.8 Å³ in the mutant. To further assess the sequence variability and structural consequences of the mutated Ω-loop, we redesigned the three substituted positions using ProteinMPNN^40^ and modelled the resulting variants using AlphaFold.^45^ Notably, the predicted Ω-loop volume of the redesigned variants (1358.8 – 1375.0 Å³) is closer to the wild-type structure than to the mutant, suggesting partial recovery of active-site architecture (*Tajane et al., unpublished*). In a follow-up study, these engineered variants will be examined experimentally using enzyme kinetics assays with penicillins and oxyiminocephalosporins to assess their effects on substrate specificity and catalytic activity.

## Conclusion

This study provides a systematic, structure-wide characterization of Ω-loops across 26,152 protein structures, comprising obligate homodimers, heterodimers, transient dimers, crystal dimers, and monomers. We show that Ω-loops are structural motifs present at an average of roughly two loops per Ω-loop-containing structure, with six-residue loops being most common and longer-residue loops occurring less frequently, consistent with earlier reports. Across all classes, non-hydrogen-bonded Ω-loops were more frequent than hydrogen-bonded ones, supporting that Ω-loops are nonregular, generally flexible, and solvent-exposed motifs. Nevertheless, the differences we observed between HBΩ and nHBΩ loops, including more compact, shorter intra-loop distances and reduced solvent exposure in HBΩ-loops show that the presence or absence of a terminal hydrogen bond does slightly affect the loop architecture, even Ω-loops non-regular structural class. Obligate homodimers contained the highest number of interface Ω-loops and Ω-loops pairs among all classes we studied. This finding aligns with another result: interface Ω-loops in obligate homodimers and heterodimers were more conserved than non-interface Ω-loops. As previously reported, the obligate homodimer interface is generally more conserved than in other datasets.^43^ Our analysis of amino acid preference and propensity also confirms that glycine and proline are the most common residues in Ω-loops across all datasets, in line with their role in backbone flexibility and turn formation. The limited sequence variation in the ProteinMPNN result, along with the conservation pattern from Rate4Site, indicates that Ω-loops remain nonregular yet do not readily tolerate sequence change. When we categorize Ω-loops, we find out that most of the Ω-loops fall into folding- and stability-associated categories, and very few loops perform several roles at once. Taken together, despite the non-regular structure of Ω-loops, their abundance in proteins helps provide stability and folding, alongside their known roles in ligand binding, catalysis, and molecular recognition.

## Author Approvals

All authors have seen and approved the manuscript, and it hasn’t been accepted or published elsewhere.

## Author Contributions

SD have conceived the idea. ST and SP have done all the experiments and analyses. SD and ST wrote the manuscript with input from all authors.

## Acknowledgments

ST has a fellowship from UGC-NET. SP acknowledges MHRD fellowship. SD acknowledges RLS grant from DBT India.

## Data Availability

The datasets and the data analysis codes used in the study are available at https://figshare.com/s/68cec2a2eb61c23c03ed

## References

1. Morris R, Black KA, Stollar EJ. Uncovering protein function: from classification to complexes. Essays Biochem. 2022;66(3):255–285.

2. Fetrow JS. Omega loops: nonregular secondary structures significant in protein function and stability. FASEB J. 1995;9(9):708–717.

3. Richardson, J.S. The Anatomy and Taxonomy of Protein Structure. In: Advances in Protein Chemistry. Vol 34. Academic Press; 1981:167–339.

4. Ravi Chandra B, Gowthaman R, Akhouri RR, Gupta D, Sharma A. Distribution of proline-rich (PxxP) motifs in distinct proteomes: functional and therapeutic implications for malaria and tuberculosis. Protein Eng Des Sel. 2004;17(2):175–182.

5. Kleiger G, Grothe R, Mallick P, Eisenberg D. GXXXG and AXXXA: common alpha-helical interaction motifs in proteins, particularly in extremophiles. Biochemistry. 2002;41(19):5990–5997.

6. Tajane SV, Thakur A, Acharya S, Chakrabarti P, Dey S. On the abundance and importance of AXXXA sequence motifs in globular proteins and their involvement in CC interaction. J Struct Biol. 2024;216(4):108129.

7. Dhar J, Chakrabarti P. O‧‧‧C═O interaction, its occurrence and implications for protein structure and folding. Proteins. 2022;90(5):1159–1169.

8. Chakrabarti P. On the pathway of the formation of secondary structures in proteins. Proteins. 2025;93(1):396–399.

9. Dhar J, Kishore R, Chakrabarti P. A novel secondary structure based on fused five-membered rings motif. Sci Rep. 2016;6:31483.

10. Rose, G. D., Gierasch, L. M. & Smith, J. A. Turns in Peptides and Proteins. In: Advances in Protein Chemistry. Vol 37. Academic Press; 1985:1–109.

11. Marcelino AMC, Gierasch LM. Roles of beta-turns in protein folding: from peptide models to protein engineering. Biopolymers. 2008;89(5):380–391.

12. Hadarovich A, Chakravarty D, Tuzikov AV, Ben-Tal N, Kundrotas PJ, Vakser IA. Structural motifs in protein cores and at protein-protein interfaces are different. Protein Sci. 2021;30(2):381–390.

13. Chou KC. Prediction and classification of alpha-turn types. Biopolymers. 1997;42(7):837–853.

14. Dhar J, Kishore R, Chakrabarti P. Delineation of a new structural motif involving NHN γ-turn. Proteins. 2020;88(3):431–439.

15. Fufezan C. The role of Buergi-Dunitz interactions in the structural stability of proteins. Proteins. 2010;78(13):2831–2838.

16. Sibanda BL, Thornton JM. Beta-hairpin families in globular proteins. Nature. 1985;316(6024):170–174.

17. Sibanda BL, Blundell TL, Thornton JM.Conformation of β-hairpins in protein structures: A systematic classification with applications to modelling by homology, electron density fitting and protein engineering. Journal of Molecular Biology. 1989;206(4):759–777.

18. Pavone V. On β-hairpin classification. International Journal of Biological Macromolecules. 1988;10(4):238–240.

19. Leszczynski JF, Rose GD. Loops in globular proteins: a novel category of secondary structure. Science. 1986;234(4778):849–855.

20. Pal M, Dasgupta S. The nature of the turn in omega loops of proteins. Proteins. 2003;51(4):591–606.

21. Papaleo E, Saladino G, Lambrughi M, Lindorff-Larsen K, Gervasio FL, Nussinov R. The Role of Protein Loops and Linkers in Conformational Dynamics and Allostery. Published online February 18, 2016. doi:10.1021/acs.chemrev.5b00623

22. Wallace CJ. Functional consequences of the excision of an omega loop, residues 40-55, from mitochondrial cytochrome c. J Biol Chem. 1987;262(35):16767–16770.

23. Replacing a Surface Loop Endows Ribonuclease A with Angiogenic Activity. Journal of Biological Chemistry. 1995;270(29):17180–17184.

24. Banerjee S, Pieper U, Kapadia G, Pannell LK, Herzberg O. Role of the omega-loop in the activity, substrate specificity, and structure of class A beta-lactamase. Biochemistry. 1998;37(10):3286–3296.

25. Stojanoski V, Chow DC, Hu L, et al. A triple mutant in the Ω-loop of TEM-1 β-lactamase changes the substrate profile via a large conformational change and an altered general base for catalysis. J Biol Chem. 2015;290(16):10382–10394.

26. Tajane SV, Pohane S, Chakrabarti P, Dey S. Abundance of Glycine-mediated O···C=O, N-H···N, and C-H···O Interactions in Homo- and Hetero-oligomeric Protein Complexes. Biochemistry. 2025;64(11):2394–2400.

27. Levy ED, Pereira-Leal JB, Chothia C, Teichmann SA. 3D complex: a structural classification of protein complexes. PLoS Comput Biol. 2006;2(11):e155.

28. Dey S, Ritchie DW, Levy ED. PDB-wide identification of biological assemblies from conserved quaternary structure geometry. Nat Methods. 2018;15(1):67–72.

29. Berman HM, Battistuz T, Bhat TN, et al. The Protein Data Bank. Acta Crystallogr D Biol Crystallogr. 2002;58(Pt 61):899–907.

30. Schweke H, Xu Q, Tauriello G, et al. Discriminating physiological from non-physiological interfaces in structures of protein complexes: A community-wide study. Proteomics. 2023;23(17):e2200323.

31. Kabsch W, Sander C. Dictionary of protein secondary structure: pattern recognition of hydrogen-bonded and geometrical features. Biopolymers. 1983;22(12):2577–2637.

32. Hubbard, S.J. and Thornton, J.M. *“*NACCESS*”,* Computer Program. http://www.bioinf.manchester.ac.uk/naccess/

33. Bahadur RP, Chakrabarti P, Rodier F, Janin J. Dissecting subunit interfaces in homodimeric proteins. Proteins: Struct Funct Bioinf. 2003;53(3):708–719.

34. Crooks GE, Hon G, Chandonia JM, Brenner SE. WebLogo: a sequence logo generator. Genome Res. 2004;14(6):1188–1190.

35. Ganesan K, Kulandaisamy A, Binny Priya S, Gromiha MM. HuVarBase: A human variant database with comprehensive information at gene and protein levels. PLoS One. 2019;14(1):e0210475.

36. Altschul SF, Gish W, Miller W, Myers EW, Lipman DJ. Basic local alignment search tool. J Mol Biol. 1990;215(3):403–410.

37. Suzek BE, Huang H, McGarvey P, Mazumder R, Wu CH. UniRef: comprehensive and non-redundant UniProt reference clusters. Bioinformatics. 2007;23(10):1282–1288.

38. Edgar RC. MUSCLE: multiple sequence alignment with high accuracy and high throughput. Nucleic Acids Res. 2004;32(5):1792–1797.

39. Pupko T, Bell RE, Mayrose I, Glaser F, Ben-Tal N. Rate4Site: an algorithmic tool for the identification of functional regions in proteins by surface mapping of evolutionary determinants within their homologues. Bioinformatics. 2002;18 Suppl 1:S71–S77.

40. Dauparas J, Anishchenko I, Bennett N, et al. Robust deep learning-based protein sequence design using ProteinMPNN. Science. 2022;378(6615):49–56.

41. McDonald IK, Thornton JM. Satisfying hydrogen bonding potential in proteins. J Mol Biol. 1994;238(5):777–793.

42. Martin AC, Toda K, Stirk HJ, Thornton JM. Long loops in proteins. Protein Eng. 1995;8(11):1093–1101.

43. Mintseris J, Weng Z. Structure, function, and evolution of transient and obligate protein–protein interactions. Proceedings of the National Academy of Sciences. 2005;102(31):10930–10935.

44. Sondka Z, Dhir NB, Carvalho-Silva D, et al. COSMIC: a curated database of somatic variants and clinical data for cancer. Nucleic Acids Res. 2024;52(D1):D1210–D1217.

45. Jumper J, Evans R, Pritzel A, et al. Highly accurate protein structure prediction with AlphaFold. Nature. 2021;596(7873):583–589.

